# The *Haemophilus influenzae* in vivo gene expression reveals major clues about bacterial central metabolism, acquisition of trace elements, and other essential pathways during infection of the human lung

**DOI:** 10.1101/2023.03.14.532706

**Authors:** Linnea Polland, Yi Su, Magnus Paulsson

## Abstract

*Haemophilus influenzae* is a major cause of community and hospital acquired pneumonia. While extensively studied in various laboratory models, less is known about how this species persists and causes infection inside the human lung. We present the first study on the H. influenzae in vivo transcriptome during pneumonia, and contrast this with isolates cultured in vitro under standard laboratory conditions. Patients with pneumonia were recruited from emergency departments and intensive care units in a Swedish referral hospital during 2018-2020 (n=102). Duplicates of lower respiratory samples were collected for bacterial culture and RNA-extraction. Patient samples with *H. influenzae* (n=18) from which bacterial mRNA of adequate quantity and quality could be extracted (n=8) underwent RNA-sequencing, along with duplicates of lab-cultured counterparts (n=7). The transcripts were aligned to core and pan genomes created from 15 reference strains. While in vitro bacteria clustered tightly in principal component analyses of core genome (n=1067) expression, the in vivo samples displayed diverse transcriptomic signatures and did not group with their lab-grown counterparts. In total, 328 core genes were significantly differentially expressed between in vitro and in vivo conditions. The most upregulated genes in vivo included the transferrin-acquisition genes tbp1 and fbpA and the reductase gene msrAB involved in stress response pathways. Biosynthesis of nucleotides/purines, response-to-heat systems, and molybdopterin-scavenging processes were also significantly upregulated in vivo. Major metabolic pathways and iron-sequestering processes were downregulated in vivo. In conclusion, extensive transcriptomic differences were found between bacteria collected in the human lung during pneumonia and isogenic bacteria cultured in vitro.

## Introduction

*Haemophilus influenzae* is the second most common bacterial infectious agents responsible for community-acquired pneumonia and the most frequent bacterial cause of exacerbations of chronic obstructive pulmonary disease (COPD) (1, 2). The severity of pneumonia caused by *H. influenzae* ranges from mild to very severe and symptoms typically involve an acute onset of cough and fever.

*H. influenzae* has evolutionally developed into a pathogenic bacterial species that survives exclusively in the human respiratory tract. It carries a relatively small genome of about 1,800,000 bp that is well defined and several annotated reference genomes exist, including the strain Rd KW20 which was the first free-living organism to have its entire genome established by sequencing (3). Traditionally, *H. influenzae* isolates are clustered into groups based on the presence of antigens in the polysaccharide capsule (serotype a-f) or the lack thereof (Non-typeable *Haemophilus influenzae*; NTHi). The designated serotypes correspond to genomic clusters with limited genetic variation, while NTHi is a more heterogenous group that are subdivided into clades. On average, only 47% of *H. influenzae* genes are shared between all strains and belong to the core genome (4).

Despite the genetic variations, the small genome necessitates a low genetic redundancy, making *H. influenzae* attractive as a model organism for human-pathogen interaction studies. Up until now, such studies are mainly performed *in vitro* using laboratory models or animal models, which is complicated by the restricted niche of this pathogen and specific requirements for growth in laboratory environments, including nutrient- and CO_2_ requirements. Laboratory models for studies of bacterial physiology or host-pathogen interactions inevitably introduce simplifications and experimental biases. For this reason, they only partly reflect the complexity of the environment *in vivo*. Studies on the *Pseudomonas aeruginosa* transcriptome comparing various *in vitro* models to *in vivo* samples collected from the lower respiratory tract of patients with cystic fibrosis have revealed a significant discrepancy between the gene expression of bacterial cells cultured in any *in vitro* model and bacterial cells growing in the human lung (5).

This is complicated by variations *in vivo* in physiological parameters, including altered temperature and oxygen availability, as well as limitations in nutrient availability and bacterial stresses caused by host-pathogen interactions and administered drugs. The *in vivo* transcriptome of *H. influenzae* is yet unknown and our knowledge of gene expression characteristics of this bacterial species is based on *in vitro* studies. We hypothesize that there is a significant discrepancy between the transcriptomic signatures of *H. influenzae* cells cultured in a laboratory environment and cells growing in the human lung during acute pneumonia, and that this difference can be used to find biological processes with importance for *H. influenzae* pathogenesis and persistence *in vivo*. In the present study we aimed to elucidate this difference by comparing the gene expression of *H. influenzae* grown in human airways and *in vitro*.

## Methods

### Study participants and sample collection

Study participants were recruited among patients admitted to the Emergency Departments at Skåne University Hospital (Lund and Malmö, Sweden) between June 2019 and May 2020. Inclusion criteria were 18 years of age or older and two of the following signs: productive cough, dyspnea, and/or fever (≥ 38.5° C). In addition, participants were recruited among patients admitted to the intensive care unit at Skåne University Hospital (Malmö, Sweden) between July 2018 and March 2020 (Figure 1). Inclusion criteria were 18 years of age or older and clinician’s suspicion of pneumonia based on the decision to collect respiratory samples for microbiological analysis. Sputum or bronchoscopy guided bronchial wash samples were collected using 2 x 10 ml sterile 0.9% saline. A portion of the samples were sent for bacterial culture and RNAlater solution (Thermo Fisher Scientific, Waltham, MA, USA) was added to the remaining portion of the samples within one minute of collection to preserve RNA integrity. All samples with RNAlater were stored at −20° C. Study data were collected from the hospital records using an electronic case report form (REDCap) hosted at Lund University (6). The present project was approved by the Swedish Ethical Review Authority (Dnr 2019-01012). All patients were given oral and written information about the study and signed a consent form before analysis begun.

**FIGURE 1:**
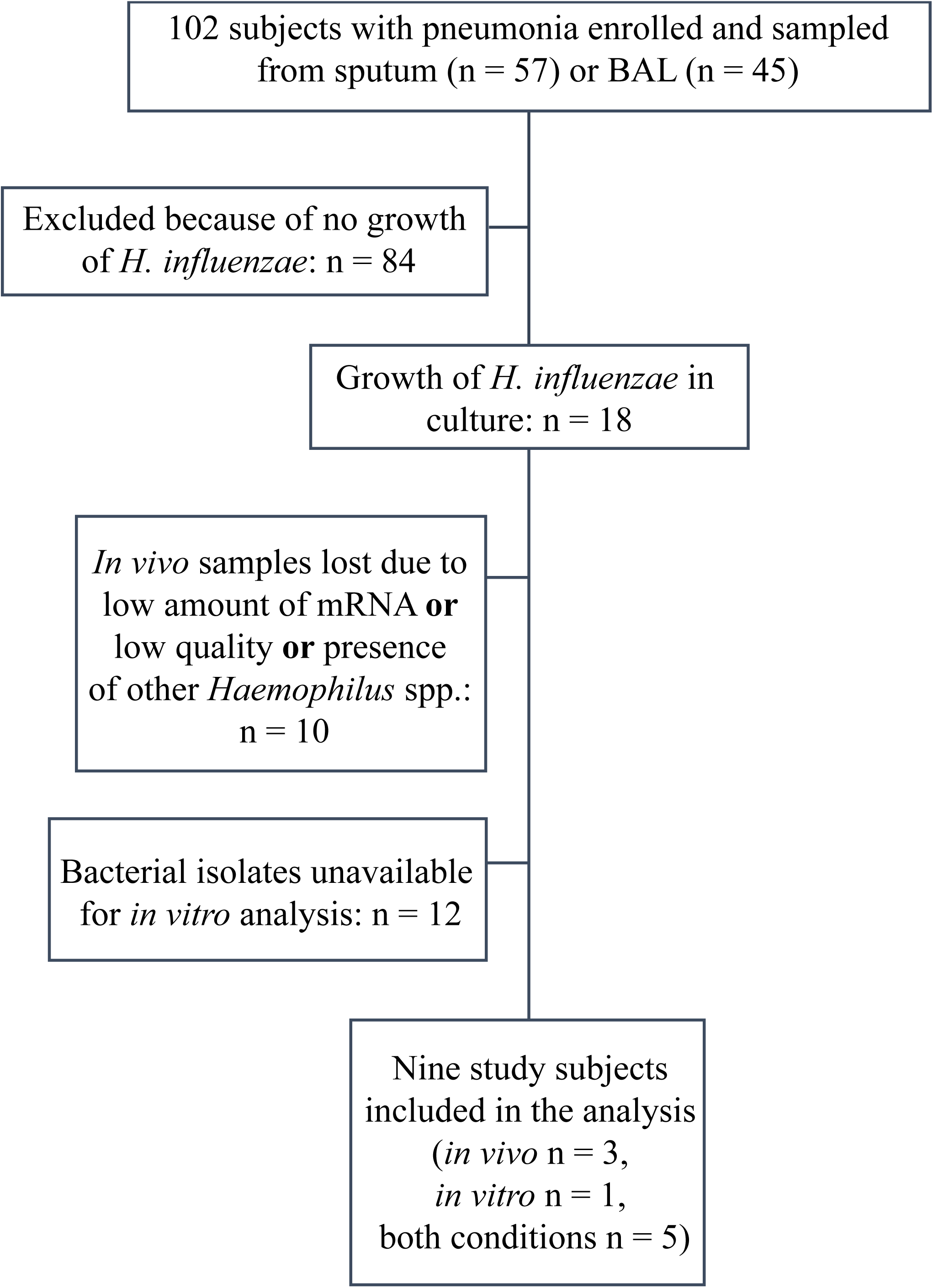
Flow chart of study subject and sample selection.

### Bacterial strains and culture conditions

Samples underwent standard bacterial culturing at Clinical Microbiology, Laboratory medicine Skåne (Lund, Sweden). Bacterial species were identified based on phenotype and MALDI-TOF (Matrix assisted laser desorption/ionization – time of flight), and later confirmed with taxonomic sequence analysis using Kraken2 (7). Bacterial isolates were put in glycerol stock and frozen at −80° C for later analysis. The reference strain Non-typeable *H. influenzae* 3655 (8) was kindly provided by Prof. Riesbeck (Lund university, Sweden).

For *in vitro* gene expression analysis, *H. influenzae* isolates were inoculated to initial OD_600 nm_ of 0.05 in brain heart infusion (BHI) broth with addition of 10 μg/ml nicotinamide adenine dinucleotide (NAD) and 10 μg/ml hemin. The isolates were cultured at 37°C, in 5% CO_2_ and 200 rpm shaking and stopped in exponential phase after 240 minutes with 2 minutes of centrifugation at 10,000 x g to obtain a pellet. The supernatant was removed and RNAlater was added to the samples.

### RNA extraction, rRNA depletion, library preparation and sequencing

RNA was extracted from *in vitro* bacterial suspensions or clinical samples in RNAlater solution using a phenol-chloroform based protocol (Trizol, Thermo Fisher Scientific). For a detailed description of this process, and of the depletion of ribosomal RNA (rRNA), library preparation, and sequencing procedure, we refer to Supplement File 1.

### Creation of core and pan genome

The core and pan genome of the 15 *H. influenzae* reference genomes listed in the KEGG database (Supplement Table S1) were created with Roary/15.36.42 and annotated with Prokka/1.11 (9, 10). This annotation was manually complemented by running BLAST on the nucleotide sequence and adding (primarily) gene names or (secondary) loci from *H. influenzae* Rd KW20 when available. If the gene was not in the Rd KW20 genome, *H. influenzae* PittII loci were used. Some genes in the pan genome were not possible to identify and were assigned the locus tag prefix “group”. The core genome was defined as genes present in all 15 reference strains and the pan genome as the full list of genes present in one or more of these strains.

### Bioinformatic data analysis

Initial computations were performed at the high-performance computing cluster Uppsala Multidisciplinary Center for Advanced Computational Science (UPPMAX) GDPR-compliant server Bianca. Raw reads of the transcripts were processed and a multi locus sequence type (MLST) were assigned to each isolate as described in Supplement File 1. The DESeq2/1.32.0 package (11, 12) in R/4.2.0 statistical programming language (13) was used for subsequent differential gene expression and statistical analysis. Genes with less than 10 counts in total were filtered out. *P* values and adjusted *P* values were calculated with Wald test and adjusted with Benjamini-Hochberg to assess the significance of differences in mean expression between *in vivo* (clinical) and *in vitro* (lab-cultured) isolates. Differentially expressed genes (DEGs) were defined as genes showing a statistically significant (adjusted *P* value <0.05) log2 fold change (logFC) of ≥1 between bacteria grown *in vivo* and *in vitro*.

Plots were created in R/4.2.0. Principal component analyses (PCA) were performed after variance stabilizing transformation. Clustered heatmaps were produced using Pearson correlation and Ward’s method on variance stabilized counts.

### Gene ontology and subsequent statistical analysis

Gene ontology (GO) classification of core genes was performed using the PANTHER classification system, version 16 (14). Further analysis of statistical overrepresentation and enrichment was done using tools provided in the topGO package in R/4.2.0 (15). When calculating statistical fold enrichment of GO terms, the DEGs were divided into up- and downregulated genes, comparing the respective groups to the rest of the core genome. The weight01 algorithm and Fisher statistics were used for this analysis. The weight01 algorithm is a mixture of the elim and weight algorithms that takes GO hierarchies into account when calculating *P* values, leading to a more conservative result when compared to the classic algorithm (16). As this method accounts for the GO topology the tests are not independent, rendering corrective measures for multiple testing inapplicable.

### Data availability

FASTQ files depleted of human genomic reads are available from the European Nucleotide Archive with study accession number PRJEB60515. All R code used in this study can be made available upon reasonable request.

## Results

### Study participants, *in vivo* and *in vitro* samples

Out of the 102 participants recruited in the study (N = 57 and N = 45 from the ER and the ICU, respectively), lower respiratory tract samples from 18 participants contained culturable *H. influenzae* (Figure 1). RNA was extracted from these and sequenced at between 215 and 509 M reads per sample (mean 341.5 M reads). Resulting reads were taxonomically classified and samples that did not pass quality control or that contained other species in the *Haemophilus* genus were excluded. Eight samples were ultimately deemed appropriate for downstream bioinformatic analysis. In addition, we picked two colonies of *H. influenzae* from the routine cultures at the clinical microbiology laboratory from six participants and one reference strain (*H. influenzae* 3655); these were cultured in broth under routine conditions, and RNA was then extracted and sequenced with 15.7-22.0 M reads (average 18.5 M reads) (Figure 1).

Clinical and demographical data of the study cohort is presented in Table 1, with a full description of each study participant in Supplement Table S2. MLST genotypes and phenotypic traits of included bacterial strains are found in Table 2, and basic sequencing data in Supplement Table S3.

**Table 1.** Demographic data of included study subjects Demographics and clinical data of study cohort. Values indicate N (%) unless otherwise specified. Comorbidities, previous and ongoing medications, vital sign assessment, and laboratory values were retrieved from the hospital records at the emergency departments (ED) or intensive care units (ICU), and qSOFA-scores were calculated from these values. No study subject suffered from cystic fibrosis, bronchiechtasis, pulmonary fibrosis, pulmonary cancer, or other tumor disease. Smoking history includes both current (N = 3) and previous (N = 3) smoking habits; all patients admitted to the ICU were current smokers. Abbreviations: COPD; chronic obstructive pulmonary disease, qSOFA; quick Sepsis Organ Related Failure Assessment.

| Characteristic | Study population<br>(*N* = 9) |
| --- | --- |
| Mean age (SD), yr | 67.9 (12.3) |
| Sex. M | 5 (56) |
| Previous medical conditions and comorbidities |  |
| *COPD/Asthma/Emphysema* | 6 (67) |
| *Diabetes mellitus* | 2 (22) |
| *Immunodeficiency* | 3 (33) |
| *Smoking history* | 6 (67) |
| Antibiotic treatment last 5 days | 3 (33) |
| Clinical data at enrollment |  |
| *Purulent sputum* | 6 (67) |
| *Temp > 38C or < 35C last 24 h* | 5 (56) |
| *New pulmonary infiltrate* | 7 (78) |
| *Mean pneumonia-symptom duration (SD), days* | 3.2 (3.5) |
| *Median qSOFA total points (range)* | 1 (0-3) |
| *Mean plasma-CRP (SD), mg/l* | 211 (123.9) |
| *Mean white blood cell count (SD), 10<sup>9</sup>/l* | 13.9 (3.6) |
| Admission type |  |
| *Not admitted* | 1 (11) |
| *Ward* | 5 (56) |
| *ICU* | 3 (33) |
| ICU outcomes |  |
| *Mean FiO2 (SD), %* | 53 (21) |
| *Mean length of ventilator treatment (SD), days* | 1.3 (0.6) |
| 30-day mortality | 2 (22) |

**Table 2.** Microbiology data of all samples Microbiologic characteristics of each clinical sample with phenotypic resistance as detected by routine clinical susceptibility testing. No isolate harbored chromosomal resistance, but decreased susceptibility to trimethoprim-sulfamethoxazole (TSU) was commonly reported. The culture results indicated that three samples contained additional bacterial species to Haemophilus influenzae, including Moraxella catarrhalis (M.cat), Streptococcus pneumoniae (Pnc), Staphylococcus aureus (S.aureus), Klebsiella aerogenes (K.aer), Candida albicans (C.alb), and a mixed microbial flora (mxfl). The percentage of transcripts aligning to the H. influenzae core genome in vivo sample with Kallisto were above 1% for all included samples. When possible, transcripts were de novo assembled and assigned to multi locus sequencing type (MLST) groups, but for seven isolates this was not possible. These did not belong to previously described ST-types (most often differing by gene fucK) and are presented as “new”. Note that LUIN_26 contained two distinct H. influenzae strains.

| Study ID | LUIN_26 | LUIN_28 | LUIN_29 | LUIN_31 | LUIN_33 | MAAK_03 | MAIV_24 | MAIV_32 | MAIV_34 |
| --- | --- | --- | --- | --- | --- | --- | --- | --- | --- |
| Bacteria in sample other than *H. influenzae* | No | No | No | No | M.cat | No | Pnc, S.aureus | No | Pnc, K.aer, C.alb, mxfl |
| Betalactamase detected | No | No | No | Yes | Yes | No | No | No | No |
| Other phenotypic resistance | No | No | TSU | TSU | No | No | No | TSU | TSU |
| MLST | ST-12 and new | ST-266 | New | New | New | New | ST-12 | New | New |

All downstream bioinformatic analyses aim to compare *in vivo* to *in vitro* cultured bacteria.

### Core and pan genome

The pan and core genomes were created using a convenience selection of 15 of the most commonly used lab and reference *H. influenzae* strains, including genomes from NTHi serotype A, B, D, and F. The resulting core and pan genomes comprised 1072 and 4163 genes, respectively. A large proportion of the identified genes belong to the “cloud genome”, i. e. collection of genes that were present in only one of the included reference genomes (1942 genes, 46.6%; Supplement Figure S1A-C). 1067 of the core and 3200 of the pan genes were found in the 9 isolates and reference strain NTHi3655 included in the present study.

### Differentially expressed genes between bacteria *in vivo* and *in vitro*

To identify genes that were expressed differently between *H. influenzae* growing in the lung and in standard laboratory conditions, an unguided differential expression analysis was performed. In the 328 genes of the core genome defined as DEGs in this process (30.7%), 130 were upregulated (39.6%) and 198 (60.4%) were downregulated *in vivo*. In total, 489 of core genome genes were upregulated *in vivo* (45.8%), while 578 were downregulated (54.2%) (gene expression lists are presented in Supplement File 2, 3, and 4).

### Unsupervised clustering and data visualization

For dimensionality reduction and visualization, principal component analyses (PCA) were performed including all core (Figure 2A) and pan genome genes (Figure 2B). In the core genome analysis, the *in vitro* cultured bacterial isolates cluster closely together regardless of genetic background. The *in vivo* gene expression showed a higher transcriptional diversity and did not cluster with the corresponding bacterial isolate cultured *in vitro*. In the pan genome PCA the *in vitro* gene expression is less homogenous and isolates form a less tight cluster.

**FIGURE 2:**
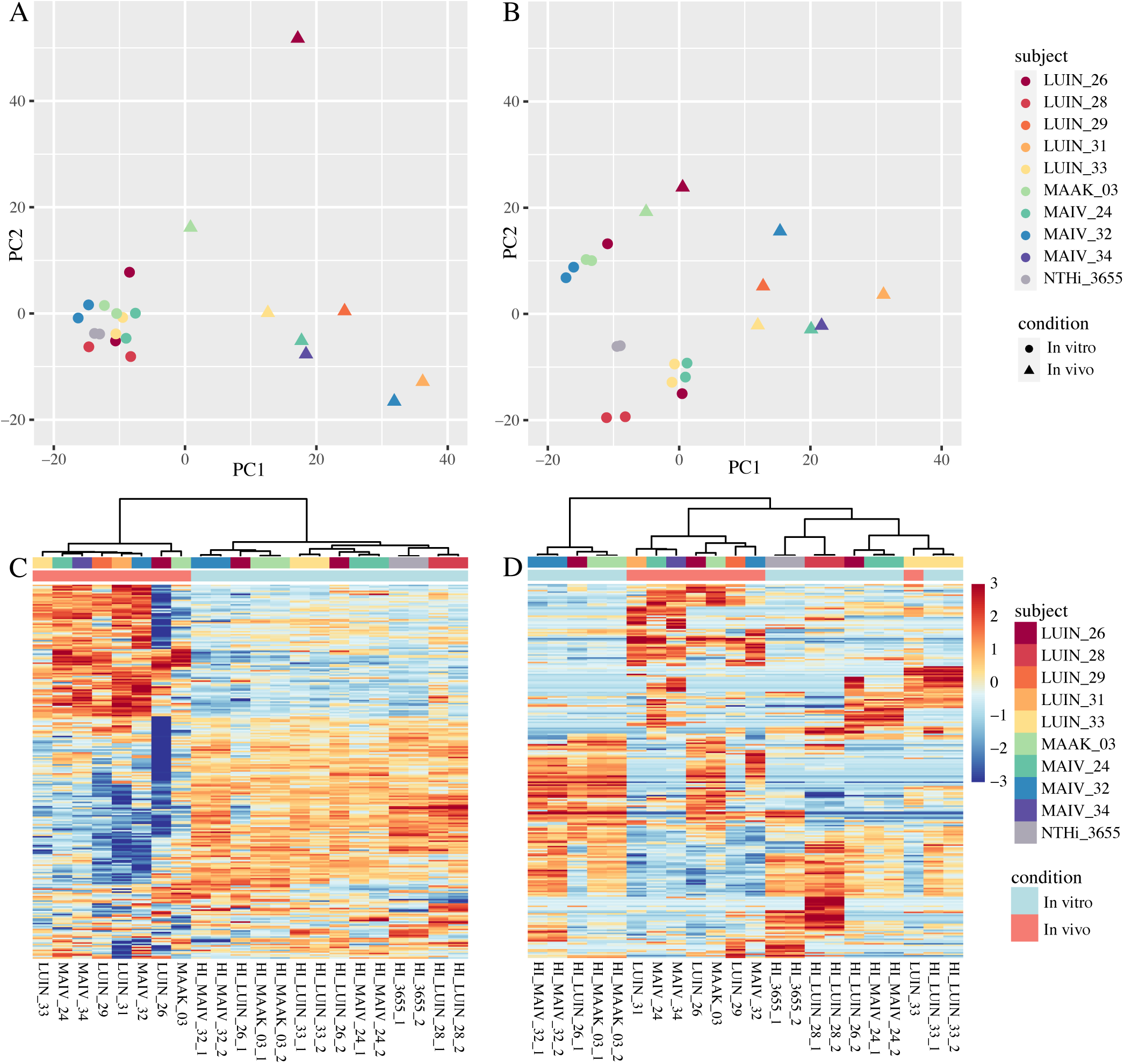
PCA-plot of expression of all core (A) and pan genes (B) and heatmaps displaying the normalized counts of the top 250 most variable genes included in the core (C) and pan genome (D). In the PCA plot *in vitro* isolates are represented by dots and *in vivo* isolates by triangles, each color representing one study subject. In the core genome analysis, *in vitro* isolates clusters tightly together, while *in vivo* samples show more heterogeneity in their transcriptional profiles (A, PC1: 32% explained, PC2: 20% explained). In the pan genome analysis this pattern is less obvious but still apparent, although *in vitro* isolates seem to cluster in two groups rather than one (B, PC1: 22% explained, PC2: 19% explained). In the heatmap of top variable core genome genes (C), using unsupervised clustering, the *in vitro* and the *in vivo* isolates cluster with their respective culture condition rather than with the corresponding isolate. Genes of *in vivo* isolates also show more diversity with higher contrast in this analysis, while the transcriptome of *in vitro* isolates is more homogenic in appearance. The legend scale indicates the log transformed normalized counts, cropped at −3 and +3. In the top 250 variable pan genome genes, clustering is more affected by the genetic background of the isolates. Plots created in R using the DESeq2, pheatmap, and ggplot2 package.

The same pattern of clustering by growth condition is seen in heatmap displaying the logFC of the top variable core genome genes, where clinical isolates show more heterogeneity with higher contrast compared to the more homogenic appearance of lab-grown samples (Figure 2C). In the pan genome heatmap (Figure 2D) this pattern is less clear, and one *in vivo* sample (LUIN_33) instead clusters with its lab-cultured counterpart.

In general, all analyses based on the pan genome displayed high heterogeneity, probably due to the many genes present in only one or few isolate(s) (demonstrated by almost half of the genes in the pan genome belonging to the cloud genome). Since a high expression of genes present in only one or few strains would get an unproportionally large statistical bias, subsequent analyses focused on the core genome.

### Top differentially expressed genes

Despite large variations in host and clinical parameters, many genes exhibited large difference in expression levels between groups in combination with stable within-group expression. This is visualized by a volcano plot of core genes (Figure 3) and a heatmap in which we selected the 30 core genome genes with the highest absolute logFC changes between the two conditions (Figure 4). The corresponding figures for the pan genome are shown in Supplement Figure S2 and S3.

**FIGURE 3:**
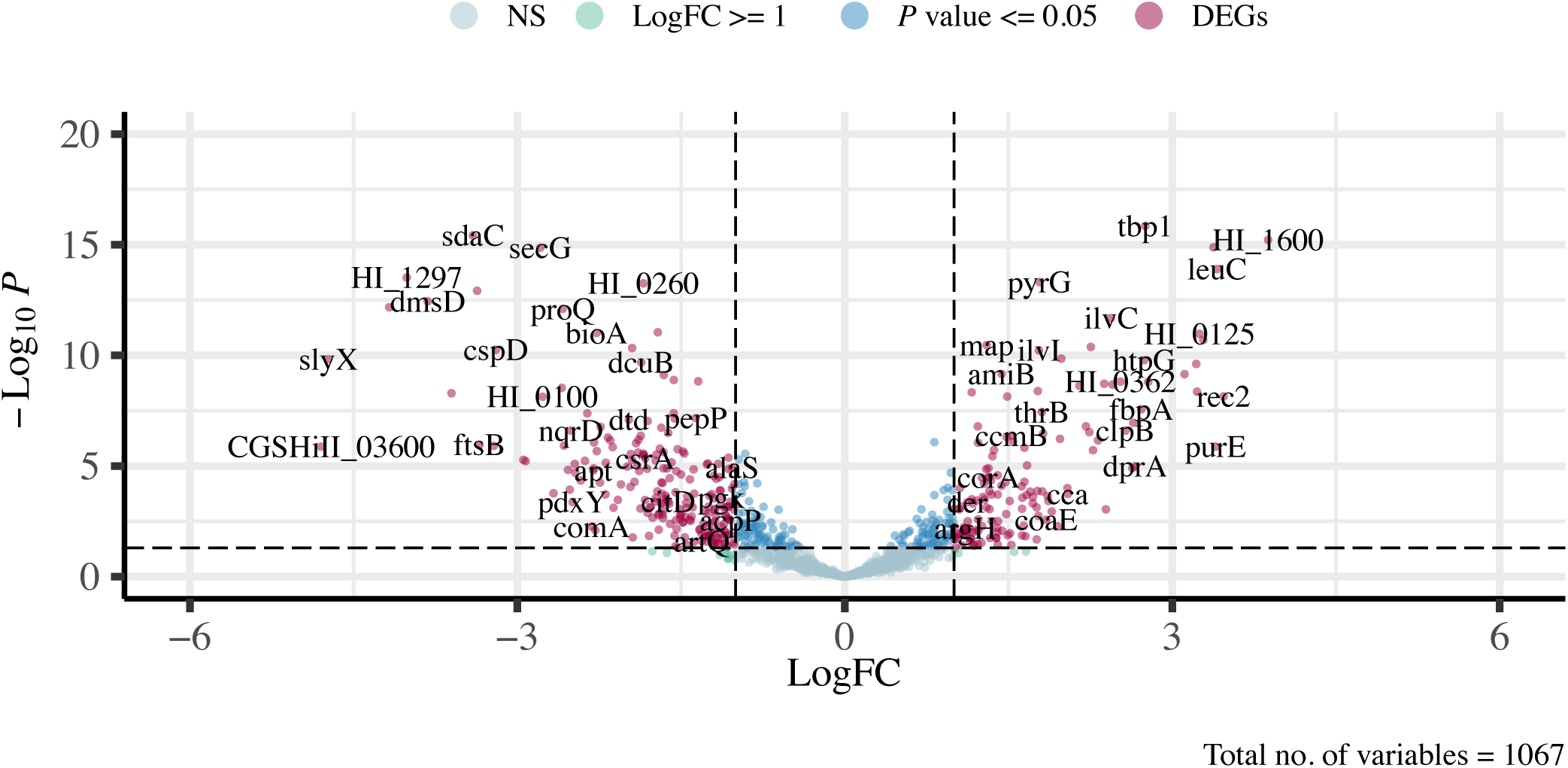
Volcano plot displaying the dispersion of core genes separated by logFC (x-axis) and p-value (y-axis, log_10_-transformed and inverted adjusted p-values). Selected gene names and loci of the most differentially expressed genes shown in graph.

**FIGURE 4:**
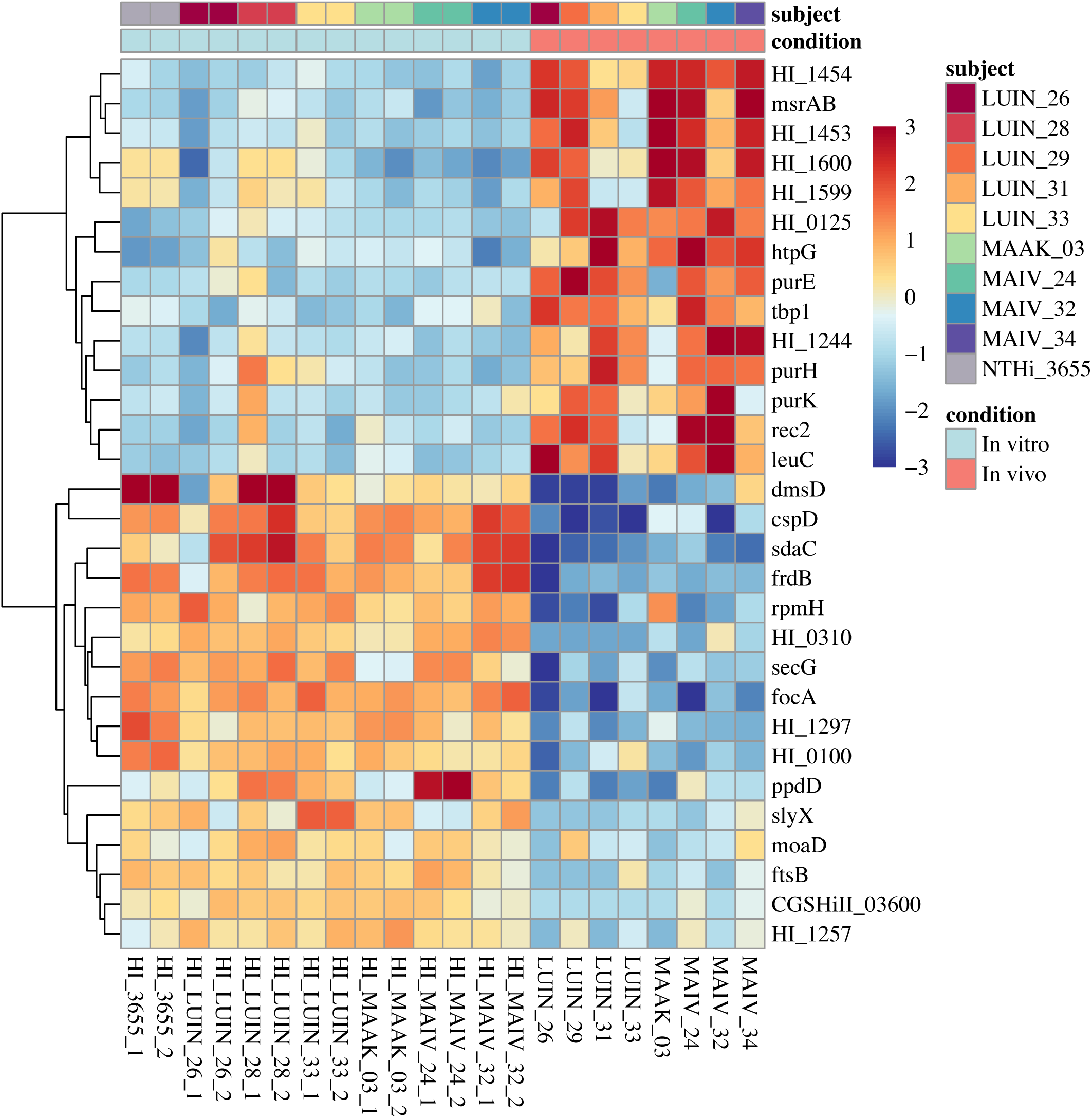
Heatmap displaying the top 30 differentially expressed genes, i.e. showing the highest absolute logFC. 14 genes showed higher (red) and 16 lower (blue) expression *in vivo*. The legend scale indicates the log transformed normalized counts, cropped at −3 and +3.

Several of the top core DEGs with higher expression *in vivo* were involved in similar pathways or related by analogous predicted functions. For instance, the gene products of *HI_1453* and *HI_1454*, with largely unknown functions, were predicted to interact with *msrAB*, involved in the response to oxidative stress. Similarly connected are the genes *purE*, *purH* and *purK* (inosine monophosphate/IMP *de novo* biosynthesis), *rec2* and *dprA* (probably involved in transformation), *tbp1* and *fbpA* (import of transferrin and iron, respectively) and *HI_1599* and *HI_1600* (both with unknown function) (14, 17).

### Enrichment of gene ontology terms and further analyses of biological processes

Gene ontology annotations of all DEGs were retrieved from the PANTHER database and the TopGO package in R. “Biological process” was determined to be the most relevant classification to explore further and use as a comparison between the DEGs and the rest of the core genome.

Of the significantly enriched GO terms, 11 were found in the downregulated and 13 in the upregulated DEGs. All significantly enriched biological process terms are presented in Table 3 and 4. A list of the 50 most enriched biological processes for the down- and upregulated DEGs is available in Supplement Table S4 and S5.

**Table 3.** Top Biological process, downregulated genes Top Biological process significant GO terms enriched for the downregulated DEGs. Processes involved in transport and sequestering of iron and cell metabolism present in the top downregulated terms (i.e. enriched in the downregulated DEGs). “Annotated” gives the total number (n) of genes present in core genome annotated to each term. “Significant” gives the number of these genes belonging to the DEGs, while “Expected” gives the expected number of annotated genes present in DEGs (i.e. same proportion as in core genome). Fold enrichment equals “Significant” divided by “Expected”.

| Gene ontology ID | Term | Annotated (n) | Significant (n) | Expected (n) | Fold enrichment | Fisher's exact test |
| --- | --- | --- | --- | --- | --- | --- |
| GO:0006108 | malate metabolic process | 3 | 3 | 0.5 | 5.66 | 0.005 |
| GO:0006094 | gluconeogenesis | 7 | 4 | 1.2 | 3.25 | 0.020 |
| GO:0006826 | iron ion transport | 8 | 4 | 1.4 | 2.86 | 0.030 |
| GO:0046854 | phosphatidylinositol phosphate biosynthetic process | 2 | 2 | 0.3 | 5.71 | 0.030 |
| GO:0046855 | inositol phosphate dephosphorylation | 2 | 2 | 0.3 | 5.71 | 0.030 |
| GO:0019646 | aerobic electron transport chain | 2 | 2 | 0.3 | 5.71 | 0.030 |
| GO:0006880 | intracellular sequestering of iron ion | 2 | 2 | 0.3 | 5.71 | 0.030 |
| GO:0006106 | fumarate metabolic process | 2 | 2 | 0.3 | 5.71 | 0.030 |
| GO:0006654 | phosphatidic acid biosynthetic process | 2 | 2 | 0.3 | 5.71 | 0.030 |
| GO:0006099 | tricarboxylic acid cycle | 8 | 4 | 1.4 | 2.86 | 0.035 |
| GO:0006096 | glycolytic process | 13 | 6 | 2.3 | 2.63 | 0.041 |

**Table 4.** Top Biological process, upregulated genes Top Biological process significant GO terms enriched for the upregulated DEGs. Biosynthetic processes of nucleotides, nucleosides, purines and branched-chain amino acids, response to heat pathways, and molybdate transport included in the top upregulated terms (i.e. enriched in the upregulated DEGs). See legend for Table 3 for explanation of terms used in table.

| Gene ontology ID | Term | Annotated (n) | Significant (n) | Expected (n) | Fold enrichment | Fisher's exact test |
| --- | --- | --- | --- | --- | --- | --- |
| GO:0006189 | 'de novo' IMP biosynthetic process | 8 | 7 | 1 | 7.22 | 0.000 |
| GO:0009098 | leucine biosynthetic process | 5 | 4 | 0.6 | 6.56 | 0.001 |
| GO:0032784 | regulation of DNA-templated transcription, elongation | 3 | 3 | 0.4 | 8.11 | 0.002 |
| GO:0015689 | molybdate ion transport | 4 | 3 | 0.5 | 6.12 | 0.006 |
| GO:0009113 | purine nucleobase biosynthetic process | 4 | 3 | 0.5 | 6.12 | 0.014 |
| GO:0009099 | valine biosynthetic process | 5 | 3 | 0.6 | 4.92 | 0.014 |
| GO:0044210 | 'de novo' CTP biosynthetic process | 2 | 2 | 0.2 | 8.33 | 0.014 |
| GO:0032508 | DNA duplex unwinding | 9 | 4 | 1.1 | 3.64 | 0.026 |
| GO:0006164 | purine nucleotide biosynthetic process | 35 | 11 | 4.3 | 2.58 | 0.036 |
| GO:0009116 | nucleoside metabolic process | 15 | 5 | 1.8 | 2.73 | 0.039 |
| GO:0009408 | response to heat | 5 | 3 | 0.6 | 4.92 | 0.039 |
| GO:0035435 | phosphate ion transmembrane transport | 3 | 2 | 0.4 | 5.41 | 0.040 |
| GO:0009097 | isoleucine biosynthetic process | 7 | 3 | 0.8 | 3.53 | 0.042 |

The biological processes enriched in the downregulated DEGs mainly related to *H. influenzae* energy production and metabolic pathways. The processes of glycolysis (1/11), the tricarboxylic acid (TCA) cycle (3/11, including the metabolism of fumarate and malate, intermediates of the TCA cycle), and gluconeogenesis (1/11) were all less common *in vivo*. We also examined the logFCs of the 23 genes in our core genome implicated in the respiratory chain, of which all but three showed lower relative expression *in vivo* and 16 were downregulated DEGs. Three of these respiratory chain enzymes (DmsA dimethylsulfoxide /DMSO reductase, nitrite reductase subunits NrfA-C, and five of six NADH dehydrogenase subunits NqrA-E), described to be especially active in anaerobic conditions (18), belonged to the downregulated DEGs. The genes of the TCA cycle, glycolytic process, and pentose-phosphate-shunt of *H. influenzae* are shown in schematics in Figure 5, and the absolute expression of genes involved in central carbon metabolism and respiration is shown in Figure 6. Figure 7A features the normalized expression of all genes assigned to the TCA cycle.

**FIGURE 5:**
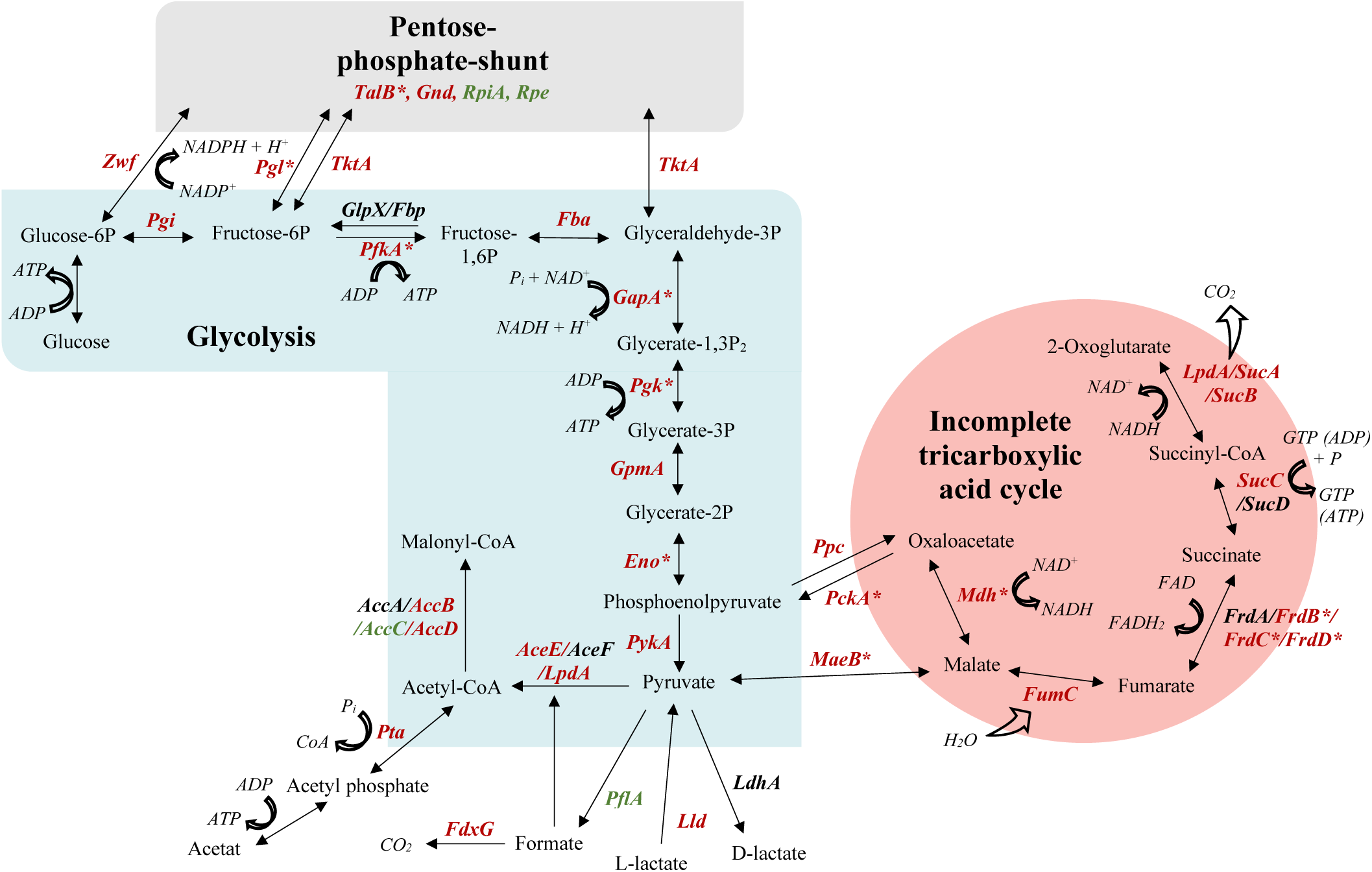
Schematics of genes and reactions involved in the tricarboxylic acid cycle, glycolysis, and pentose phosphate pathway of *H. influenzae*. Color of gene-names indicate direction of gene expression in our material (green: upregulated *in vivo*, red: downregulated *in vivo*, black: not present in core genome). *DEGs: significant (*P*<0.05) logFC >1. Based on data from López-López et al 2020 (33).

**FIGURE 6:**
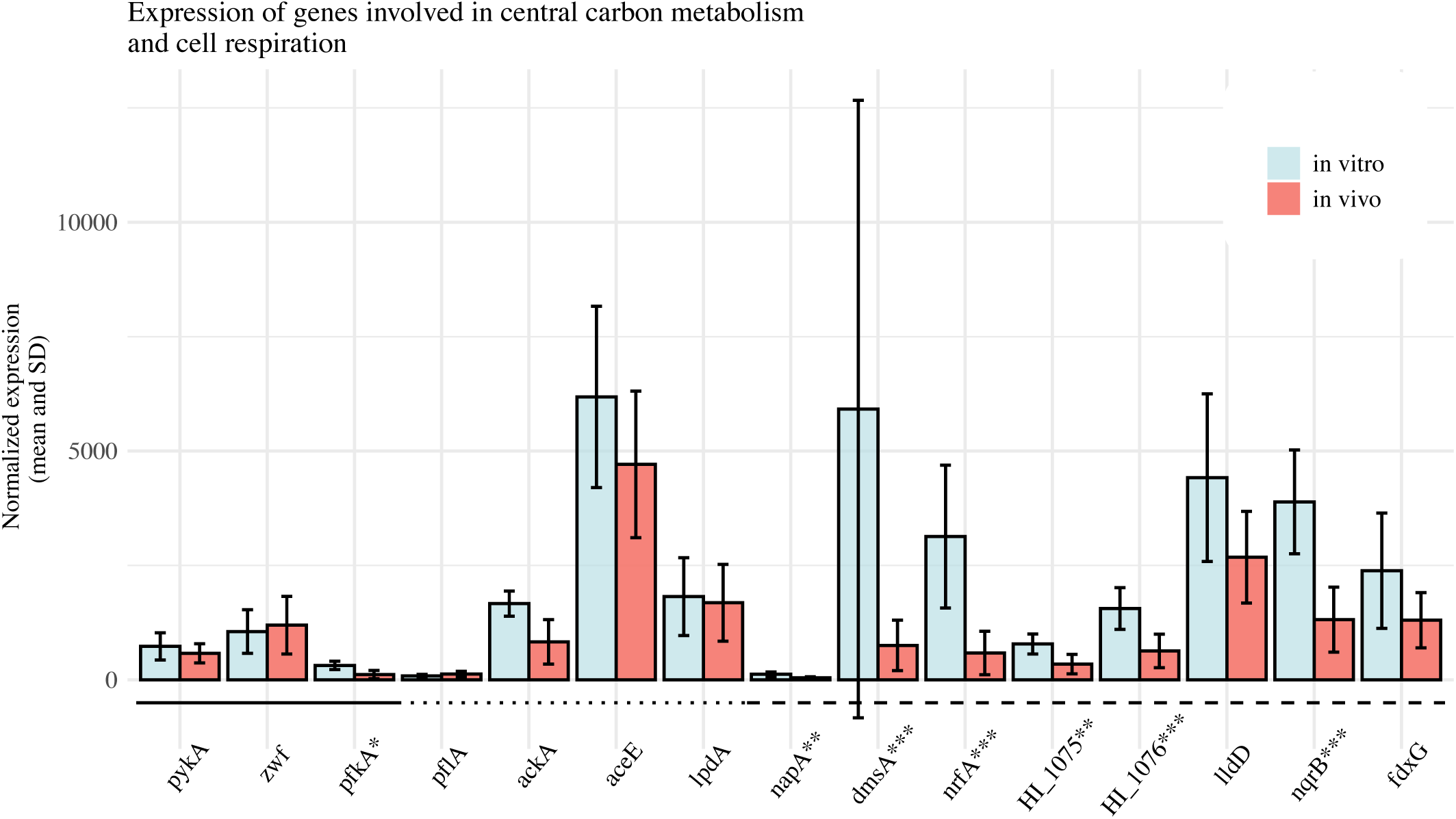
Display of the normalized expression (mean and standard deviation) of genes involved in glucose catabolism (solid line), pyruvate conversion (dotted line) and respiratory metabolism (dashed line), separated on *in vivo* and *in vitro* isolates. As discussed in the text, main difference between conditions observed for genes relevant for respiratory metabolism (e.g. *dmsA*, *nrfA*, *HI_1076, nqrB*): all included genes in this subgroup displaying higher mean expression *in vitro*. Genes included in DEGs; \*\*\**P*<0.001,\*\**P*<0.01,\**P* ≤ 0.05.

**FIGURE 7:**
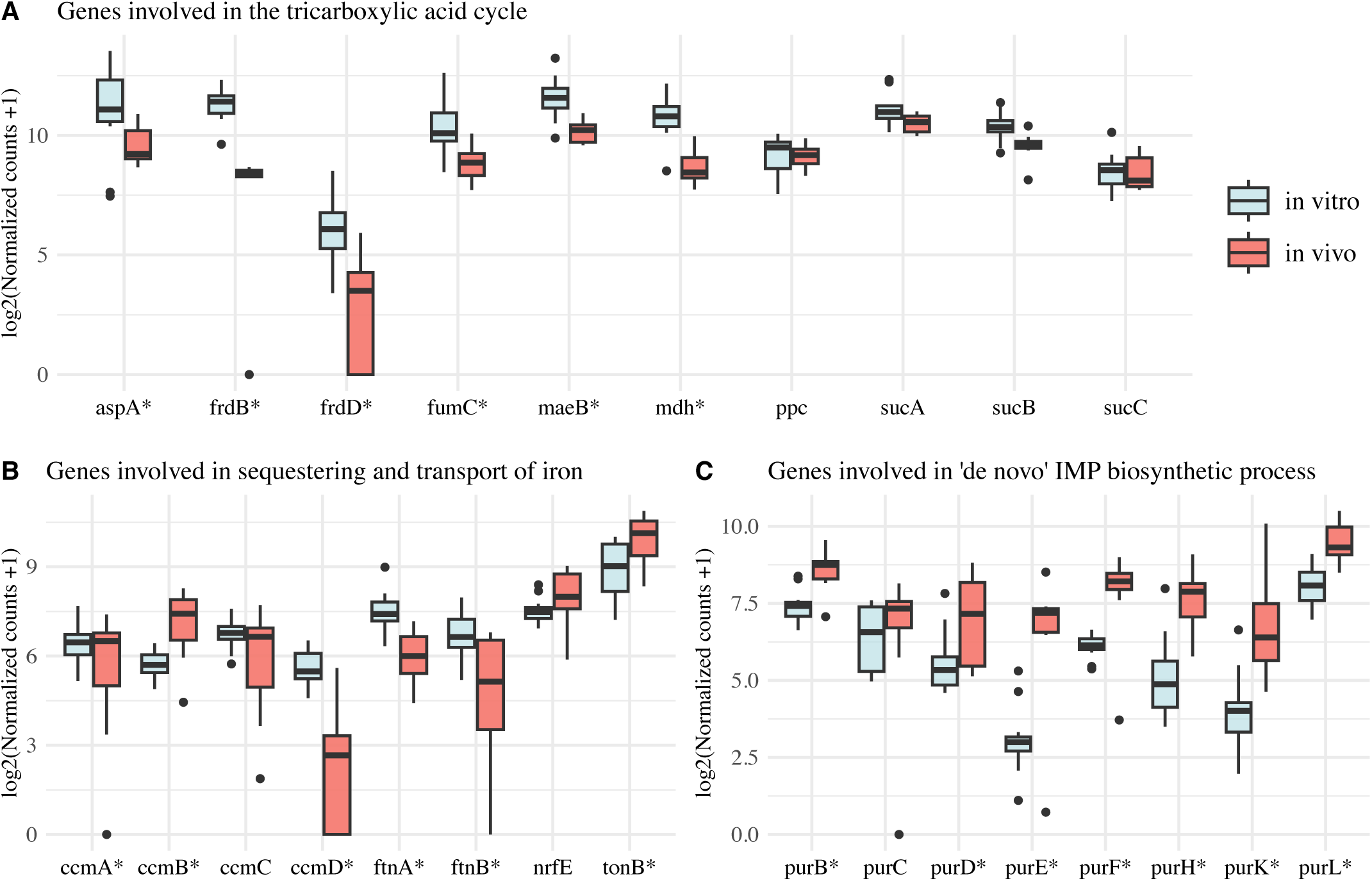
The expression of all core genes (log2 normalized counts +1) belonging to the following biological processes: significant processes relevant for the TCA cycle (A: enriched in the downregulated DEGs), iron sequestering and transport (B: enriched in the downregulated DEGs), and ‘de novo’ IMP biosynthetic process (C: enriched in the upregulated DEGs).

Iron transport and intracellular sequestering (2/11) were also less common *in vivo* compared to *in vitro* (Figure 7B). We found no other iron-related GO-terms to differ significantly between the DEGs and the rest of the core genome, including iron homeostasis (GO:0055072 or GO:0006879). Further results on the expressional changes of iron-heme related genes is found in Supplement File 5 (Iron-heme related genes).

The biological processes more frequently found *in vivo* related to biosynthesis of purines/pyrimidines and other components of DNA and RNA (5/13), biosynthesis of branched-chain amino acids leucine, valine and isoleucine (3/13), DNA unwinding and transcription (2/13), response to heat (1/13), phosphate transmembrane transport (1/13) and transport of molybdate (Mo) (1/13). The *de novo* biosynthetic process of IMP was the most enriched GO-term overall, with a majority of purine regulon genes (19) being upregulated DEGs (Figure 7C). Other than response to heat, no significant difference between the genes in the categories stress response including oxidative stress, response to different chemicals, or response to DNA damage.

The observed change in molybdate-transport resulted from all *modABC* operon genes, responsible for the influx of Mo into the bacterial cell (20, 21), being significantly upregulated *in vivo* (logFC 2.32, 1.67, 1.98, respectively). The *modE* gene was the only gene in this operon not belonging to the positive DEGs, consistent with this protein functioning as a transcriptional regulator by acting as a Mo dependent repressor of this pathway (21–23). Mo-molybdopterin cofactor biosynthetic process was not included in the significant GO-terms, but many enzymes taking part in this pathway were instead less expressed *in vivo* (seven out of nine genes having a negative logFC, ranging from −2.95 to −0.47), one of which was included in the top 16 genes showing the highest decrease in expression (*moaD*).

## Discussion

The primary aim of the present study was to describe the *H. influenzae* transcriptome *in vivo* in relation to bacterial cells cultured under typical laboratory conditions. In this first clinical *in vivo* transcriptome-sequencing study of *H. influenzae* gene expression, we found several interesting themes worth exploring further.

To enable meaningful statistical transcriptomic analysis, a core genome was created comprising 1072 genes that were present in 15 reference strains, a slightly lower number than previously reported *H. influenzae* core genomes (4). This difference is likely to be explained by different methodologies and the choice of strains that went into the analysis. Many genes related to antibiotic resistance and virulence were not shared between all strains and therefore not included in the core genome. Hence, the core genome is weighted towards housekeeping genes required for basic cellular functions. In our material, 30.7% of core genome genes were differentially expressed (according to our definition of DEGs). This was lower than a previous report comparing the effect on gene expression caused by *in vitro* host-bacterial cell interactions, in which 1068 of the 1877 *H. influenzae* genes (56.9%) were differentially expressed (24).

Most notable was the observation that bacteria growing in the human lung have a distinct, but variable, transcriptomic signature compared to bacteria growing *in vitro* under controlled laboratory conditions. Despite the large variations in host factors that affect growth conditions *in vivo* compared to the homogenic culture procedures *in vitro*, bacterial cells cluster based on these environmental conditions rather than genetic background. The importance of the growth environment for the transcriptomic signature being greater than that of the genetic background was also reported by Aziz *et al.* (25). These authors reported that the transcriptomic differences for *in vitro* cultured bacteria caused by genetic adaptation to the lung was very modest in NTHi isolates collected from the nasopharynx and the lower respiratory airways. Similarly, genetically diverse *P. aeruginosa* isolates collected from human lung exhibited similar gene expression when cultured *in vitro* (26).

The gene ontology-based analysis of the core genome transcripts revealed vast differences in the central metabolism between the growth conditions, most notably the tricarboxylic acid (TCA) cycle. The TCA cycle in many *Haemophilus* spp. is known to be incomplete due to the lack of aconitase, citrate synthase, isocitrate, and succinate dehydrogenase, rendering the organism prone to anaerobic respiration highly dependent on malate dehydrogenase (3, 27–29). In our analysis, several critical genes involved in the tricarboxylic acid cycle were significantly downregulated in the clinical isolates. A plausible explanation for the general downregulation of the TCA cycle *in vivo* in this species could be that the bacterial cells are in a more dormant and less metabolically active state during clinical infection. This may be related to a biofilm growth mode (30), or to alterations in the energy production based on limitations in the supply of proteins, sugars, and fats. However, we did not find any other convincing signs of biofilm activity (see Supplement File 5: Biofilm formation). Our findings are also consistent with those of Cornforth *et al.,* who reported a general decrease in the activity of the TCA cycle in *P. aeruginosa* during human infection (31).

Although the lung constitutes an oxygen-rich environment in the healthy state, factors such as alveolar exudation and airway obstruction in inflammatory processes are known to contribute to hypoxia, creating anaerobic or microaerophilic growth conditions for microbes (32). In standard laboratory conditions, *H. influenzae* is cultured in a carbon dioxide enriched (5%) aerobic environment, a procedure also used in the present study. When comparing our data to previous studies on the metabolic flexibility of *H. influenzae* when exposed to different oxygen levels (18, 29, 33–35) (see Supplement File 5: *H. influenzae* metabolism when exposed to different oxygen levels) our results indicated contradictory conclusions about the oxygen availability for microbes in the lung.

Two significant genes for the action of competence, *dprA* and *rec2*, were among the genes that had the highest relative expression *in vivo*. The natural competence in *H. influenzae* is recognized as having a nutritional function since only 10-15% of obtained exogenous DNA is used for homologous recombination in transformation (36), while most nucleic acids are probably catabolized and used in *de novo* DNA synthesis (37, 38). The fact that the expression of competence-associated genes is influenced by nutritional signals (CRP-cAMP levels, the PTS-system, and possibly also purine depletion) (37, 39–41) and that free polymeric DNA in high amounts can be found in the mucus of both healthy and diseased human respiratory airways (42, 43), additionally supports this concept. DNA-uptake is also highly increased when isolates in exponential phase are transferred from nutrient rich media to starvation media (44). It would consequently be expected that genes involved in competence would be induced in the human respiratory tract, as opposed to in supplemented BHI-broth.

Several processes primarily concerning replication, transcription, or the creation of necessary components of these pathways, were enriched *in vivo*. They constituted seven of the 13 top enriched GO terms, with *de novo* biosynthesis of purine IMP being the most enriched term. This is somewhat surprising since the isolates cultured *in vitro* were analyzed during exponential growth phase. Bacteria living in a human host would have restricted availability to pre-synthesized nucleotides and nucleobases, necessitating increased formation of fundamental components of nucleic acids *de novo*. Corresponding with this notion, we did not see a consistent increase in the expression of genes involved in nucleoside salvage, with some genes being upregulated (*hpt*, *upp*, *HI_0862*, *apaH*) and some being downregulated (*apt*, *adk*, *udp*, *udk*, *deoD*) (14, 45). Interestingly, genes involved in purine formation have been implicated as virulence determinants in a number of pathogenic bacteria in addition to *H. influenzae*, including many species of which the respiratory system is the primary niche (46–54). For example, in a study where attenuated mutant strains in a murine model were identified, many enzymes required for the synthesis of IMP were demonstrated to be essential for *H. influenzae* infection of the lung (55).

Iron availability is limited by the human host as a defense against infectious organisms through utilization of iron-sequestering proteins such as lactoferrin in epithelial cells, regulation of extracellular iron concentration, as well as other mechanisms of nutritional immunity (56). We found many genes involved in sequestering and transport of iron to be significantly more expressed in lab-grown isolates, probably because of a relative iron repletion *in vitro*. However, two transporters of iron, transferrin-binding protein TbpA (*tbp1*), and ferric binding protein FbpA, were found among the top upregulated DEGs. Many obligate human pathogens such as *H. influenzae*, *Neisseria meningitidis,* and *Neisseria gonorrhoeae*, possess the ability to scavenge transferrin-bound iron from their host with transferrin-binding proteins analogous to TbpA, as opposed to the usage of siderophores (57–59). Both FbpA, interacting with TbpA in the accessing of transferrin iron pools in functioning as a nodal point in iron transport (60), as well as TbpA, were identified as having increased expression in iron/heme-restricted surroundings by Whitby *et al*. (61). The expression of TbpA is also known to be hampered by higher levels of hemin (62), such as in supplemented BHI broth, which would explain the observed relative reduced expression *in vitro*.

The transport of molybdate, a fundamental cofactor in several important procaryotic redox enzymes (63–67), was another pathway significantly more common in our clinical isolates. Many genes partaking in Mo-molybdopterin cofactor biosynthesis were at the same time less expressed *in vivo.* Defective molybdoenzymes have been associated with reduced virulence of various pathogenic bacteria e.g. *Escherichia coli*, *P. aeruginosa*, and *Mycobacterium tuberculosis*, and for numerous species non-functional Mo-uptake systems have been shown to create phenotypes with reduced fitness or survival (63). Pathogenic bacteria probably compete with their hosts for molybdate (63) which is only found in very low concentrations in many human body compartments such as the lung (68), making effective Mo-uptake essential in these species. Our findings indicate that while the scavenge for molybdate is intensified *in vivo*, the substrate shortage limits metabolic pathways requiring this trace element.

Response to heat-systems were also more active *in vivo*. Fever-range temperatures are vital for many aspects of the immune system’s response to infections, for example driving recruitment and activation of neutrophils (69). High body temperature has also been shown to have a direct inhibitory impact on bacterial growth and LPS production (70–72), although the effect of heat on the pathogenic ability of bacteria is debated (73). Regarding other stress response pathways, we could not find any overarching patterns, merely specific genes showing large logFCs, as exemplified below.

The reduction of methionine sulfoxide (MetSO), a product of oxygenation of small molecules, is essential for bacterial pathogens to minimize oxidative stress (74). MsrAB, a MetSO reductase, was highly upregulated *in vivo* in our material (logFC 3.38, *P*<0.001; included in the top 30 most differentially expressed genes). *H. influenzae* isolates have been observed to show increased expression of the *msrAB* gene when exposed to reactive oxygen species commonly produced by its human host (75). Lack of the gene also caused diminished survival in biofilm as well as lower invasive potential *in vitro* and in a mouse model (75). Surprisingly, MsrAB activity has been shown to be highest in temperatures of 30 degrees and lower (76). Our results indicate that transcription of *msrAB* is of importance for persistence in the human respiratory system, and that is not overtly affected by temperatures of 37 degrees and higher, as would be commonly encountered in the febrile human body.

Another MetSO reducing enzyme, DMSO-reductase, has in previous studies been reported to be essential for *H. influenzae* virulence, as mutant strains lacking the *dmsA* gene showed reduced fitness compared to the wild type (77). Remarkably, we found the DMSO reductase-involved genes *dmsA* and *dmsD* to be highly downregulated *in vivo*. One probable explanation for this is that DMSO reductase is a molybdopterin-dependent enzyme and, as noted above, the access to molybdate would likely be restricted in the lung compared to in a rich growth medium.

*In vivo* gene expression research is known for methodological difficulties, primarily since mRNA is fragile and rapidly overturned. We have taken extensive measures to report the gene expression of bacteria living in the human lower respiratory tract, collecting the samples bedside to preserve the RNA. RNA was then extracted from all samples using a combination of enzymatic and mechanical cell disruption to maximize the yield. Despite this, both the yield and the quality of extracted RNA were low, causing a large number of duplicates during sequencing which needed to be bioinformatically filtered. Sputum can be assumed to contain large amounts of RNases, which may explain the low quality of the extracted RNA. We also used a conservative approach to avoid bias caused by wrongly assigned transcripts by excluding samples that contained other species in the *Haemophilus* genus. Additionally, to avoid bias caused by low transcript numbers we excluded samples with less than 1% transcripts assigned to *H. influenzae* in the taxonomic analysis. All these factors contributed to the loss of over half of the *H. influenzae-*containing *in vivo* samples.

Some aspects of our study warrant further scrutiny. For instance, the differential gene expression we found was less distinct than that of many previous studies of *in vitro* isolates or using controlled murine protocols. Undoubtedly this is due to the inherent transcriptional heterogeneity of *in vivo* growing bacteria as a consequence of uncontrolled and differing growth environments in human airways. Moreover, we limited the biological interpretations to the core genome and compared isolates on a group level. Further interesting discoveries could have been made through pan rather than core genome analyses or by a pairwise comparison between clinical strains and their lab-grown counterparts. Such analysis could include the entire chromosome and plasmids enabling analyses of, for instance, antimicrobial resistance genes. Concerning the biological interpretations, these were based on publicly available gene ontology annotations, and the quality of this data is dependent on the quality of the annotation.

## Conclusion

In summary, we found the gene expression of *Haemophilus influenzae* to differ widely between lab-cultured and clinical isolates. *In vitro* isolates conformed in their mRNA transcription profiles, while the gene expression of *in vivo* bacteria was considerably more diverse. Isolates also clustered based on growth condition rather than genetic background. Genes involved in major metabolic pathways were less expressed *in vivo,* while the synthesis of essential components of nucleic acid, molybdopterin acquisition enzymes, and transferrin-binding proteins were upregulated in this condition. The generated dataset is suitable for hypothesis generation for future studies, including the stress response gene *msrAB* and the highly expressed hypothetical proteins HI_1599 and HI_1600. These results also raise questions about the generalizability of microbiological studies performed in controlled circumstances, which are unlikely to approximate the diverse environments encountered by pathogenic bacteria in human airways. As such, findings in our material may be particularly significant, since they point toward processes that are ubiquitous for the survival and pathogenic ability of *H. influenzae* isolates *in vivo*.

## Acknowledgements

We would like to thank students and staff at the Skåne university hospital for their help with patient enrollment and sample collection. We would also like to acknowledge Prof Marvin Whitely and Dr Dan Cornforth at Georgia institute of Technology and Prof Thomas Bjarnsholt and Blaine Fritz at University of Copenhagen for introducing us to RNA extraction methods and bacterial transcriptomics.

Support by NBIS (National Bioinformatics Infrastructure Sweden) is gratefully acknowledged.

The computations were performed on resources provided by SNIC through Uppsala Multidisciplinary Center for Advanced Computational Science (UPPMAX) under Project SNIC sens2020512 and LUNARC at Lund university. The authors would like to acknowledge Clinical Genomics Lund, SciLifeLab and Center for Translational Genomics (CTG), Lund University, for providing expertise and service with sequencing and analysis. This publication made use of the PubMLST website (https://pubmlst.org/) developed by Keith Jolley (Jolley & Maiden 2010, BMC Bioinformatics, 11:595) and sited at the University of Oxford. The development of that website was funded by the Wellcome Trust.

This study was financed by Swedish governmental funding of clinical research (ALF), Swedish research council 2018-06924 and 2021-06380, Swedish society for medical research (SSMF), The Royal physiographical society, Skåne County Council’s Research and Development Foundation and the Gyllenstiernska Krapperup foundation.

## References

1. Metlay JP, Waterer GW, Long AC, Anzueto A, Brozek J, Crothers K, Cooley LA, Dean NC, Fine MJ, Flanders SA, Griffin MR, Metersky ML, Musher DM, Restrepo MI, Whitney CG. 2019. Diagnosis and Treatment of Adults with Community-acquired Pneumonia. An Official Clinical Practice Guideline of the American Thoracic Society and Infectious Diseases Society of America. Am J Respir Crit Care Med 200:e45–e67.

2. Wilkinson TMA, Aris E, Bourne S, Clarke SC, Peeters M, Pascal TG, Schoonbroodt S, Tuck AC, Kim V, Ostridge K, Staples KJ, Williams N, Williams A, Wootton S, Devaster J-M, AERIS Study Group. 2017. A prospective, observational cohort study of the seasonal dynamics of airway pathogens in the aetiology of exacerbations in COPD. Thorax 72:919–927.

3. Fleischmann RD, Adams MD, White O, Clayton RA, Kirkness EF, Kerlavage AR, Bult CJ, Tomb JF, Dougherty BA, Merrick JM, McKenney K, Sutton G, FitzHugh W, Fields C, Gocayne JD, Scott J, Shirley R, Liu LL, Glodek A, Kelley JM, Weidman JF, Phillips CA, Spriggs T, Hedblom E, Cotton MD, Utterback TR, Hanna MC, Nguyen DT, Saudek DM, Brandon RC, Fine LD, Fritchman JL, Fuhrmann JL, Geoghagen NSM, Gnehm CL, McDonald LA, Small K v, Fraser CM, Smith HO, Venter JC. 1995. Whole-genome random sequencing and assembly of Haemophilus influenzae Rd. Science 269:496–512.

4. Pinto M, González-Díaz A, Machado MP, Duarte S, Vieira L, Carriço JA, Marti S, Bajanca-Lavado MP, Gomes JP. 2019. Insights into the population structure and pan-genome of Haemophilus influenzae. Infect Genet Evol 67:126–135.

5. Cornforth DM, Diggle FL, Melvin JA, Bomberger JM, Whiteley M. 2020. Quantitative Framework for Model Evaluation in Microbiology Research Using Pseudomonas aeruginosa and Cystic Fibrosis Infection as a Test Case. mBio 11.

6. Harris PA, Taylor R, Thielke R, Payne J, Gonzalez N, Conde JG. 2009. Research electronic data capture (REDCap)--a metadata-driven methodology and workflow process for providing translational research informatics support. J Biomed Inform 42:377–81.

7. Wood DE, Lu J, Langmead B. 2019. Improved metagenomic analysis with Kraken 2. Genome Biol 20:257.

8. Musser JM, Barenkamp SJ, Granoff DM, Selander RK. 1986. Genetic relationships of serologically nontypable and serotype b strains of Haemophilus influenzae. Infect Immun 52:183–91.

9. Seemann T. 2014. Prokka: rapid prokaryotic genome annotation. Bioinformatics 30:2068–9.

10. Page AJ, Cummins CA, Hunt M, Wong VK, Reuter S, Holden MTG, Fookes M, Falush D, Keane JA, Parkhill J. 2015. Roary: rapid large-scale prokaryote pan genome analysis. Bioinformatics 31:3691–3.

11. Love MI, Huber W, Anders S. 2014. Moderated estimation of fold change and dispersion for RNA-seq data with DESeq2. Genome Biol 15:550.

12. Soneson C, Love MI, Robinson MD. 2015. Differential analyses for RNA-seq: transcript-level estimates improve gene-level inferences. F1000Res 4:1521.

13. R Core Team. 2022. R: A language and environment for statistical computing. Repository URL https://www.R-project.org/. Retrieved 15 November 2022.

14. Thomas PD, Ebert D, Muruganujan A, Mushayahama T, Albou L-P, Mi H. 2022. PANTHER: Making genome-scale phylogenetics accessible to all. Protein Sci 31:8– 22.

15. Alexa A, Rahnenfuhrer J. 2022. _topGO: Enrichment Analysis for Gene Ontology_. R package version 2.48.0.

16. Alexa A, Rahnenführer J, Lengauer T. 2006. Improved scoring of functional groups from gene expression data by decorrelating GO graph structure. Bioinformatics 22:1600–7.

17. UniProt Consortium. 2021. UniProt: the universal protein knowledgebase in 2021. Nucleic Acids Res 49:D480–D489.

18. Othman DSMP, Schirra H, McEwan AG, Kappler U. 2014. Metabolic versatility in Haemophilus influenzae: a metabolomic and genomic analysis. Front Microbiol 5:69.

19. Mironov AA, Koonin E v, Roytberg MA, Gelfand MS. 1999. Computer analysis of transcription regulatory patterns in completely sequenced bacterial genomes. Nucleic Acids Res 27:2981–9.

20. Tirado-Lee L, Lee A, Rees DC, Pinkett HW. 2011. Classification of a Haemophilus influenzae ABC transporter HI1470/71 through its cognate molybdate periplasmic binding protein, MolA. Structure 19:1701–10.

21. Grunden AM, Shanmugam KT. 1997. Molybdate transport and regulation in bacteria. Arch Microbiol 168:345–54.

22. Walkenhorst HM, Hemschemeier SK, Eichenlaub R. 1995. Molecular analysis of the molybdate uptake operon, modABCD, of Escherichia coli and modR, a regulatory gene. Microbiol Res 150:347–61.

23. McNicholas P. 1996. The Escherichia coli modE gene: effect of modE mutations on molybdate dependent modA expression. FEMS Microbiol Lett 145:117–123.

24. Baddal B, Muzzi A, Censini S, Calogero RA, Torricelli G, Guidotti S, Taddei AR, Covacci A, Pizza M, Rappuoli R, Soriani M, Pezzicoli A. 2015. Dual RNA-seq of Nontypeable Haemophilus influenzae and Host Cell Transcriptomes Reveals Novel Insights into Host-Pathogen Cross Talk. mBio 6:e01765–15.

25. Aziz A, Sarovich DS, Nosworthy E, Beissbarth J, Chang AB, Smith-Vaughan H, Price EP, Harris TM. 2019. Molecular Signatures of Non-typeable Haemophilus influenzae Lung Adaptation in Pediatric Chronic Lung Disease. Front Microbiol 10:1622.

26. Kordes A, Preusse M, Willger SD, Braubach P, Jonigk D, Haverich A, Warnecke G, Häussler S. 2019. Genetically diverse Pseudomonas aeruginosa populations display similar transcriptomic profiles in a cystic fibrosis explanted lung. Nat Commun 10:3397.

27. Tatusov RL, Mushegian AR, Bork P, Brown NP, Hayes WS, Borodovsky M, Rudd KE, Koonin E v. 1996. Metabolism and evolution of Haemophilus influenzae deduced from a whole-genome comparison with Escherichia coli. Curr Biol 6:279–91.

28. Edwards JS, Palsson BO. 1999. Systems properties of the Haemophilus influenzae Rd metabolic genotype. J Biol Chem 274:17410–6.

29. Raghunathan A, Price ND, Galperin MY, Makarova KS, Purvine S, Picone AF, Cherny T, Xie T, Reilly TJ, Munson R, Tyler RE, Akerley BJ, Smith AL, Palsson BO, Kolker E. 2004. In Silico Metabolic Model and Protein Expression of Haemophilus influenzae Strain Rd KW20 in Rich Medium. OMICS 8:25–41.

30. Post DMB, Held JM, Ketterer MR, Phillips NJ, Sahu A, Apicella MA, Gibson BW. 2014. Comparative analyses of proteins from Haemophilus influenzae biofilm and planktonic populations using metabolic labeling and mass spectrometry. BMC Microbiol 14:329.

31. Cornforth DM, Dees JL, Ibberson CB, Huse HK, Mathiesen IH, Kirketerp-Møller K, Wolcott RD, Rumbaugh KP, Bjarnsholt T, Whiteley M. 2018. Pseudomonas aeruginosa transcriptome during human infection. Proc Natl Acad Sci U S A 115:E5125–E5134.

32. Schaible B, Schaffer K, Taylor CT. 2010. Hypoxia, innate immunity and infection in the lung. Respir Physiol Neurobiol 174:235–43.

33. López-López N, Euba B, Hill J, Dhouib R, Caballero LA, Leiva J, Hosmer J, Cuesta S, Ramos-Vivas J, Díez-Martínez R, Schirra HJ, Blank LM, Kappler U, Garmendia J. 2020. Haemophilus influenzae Glucose Catabolism Leading to Production of the Immunometabolite Acetate Has a Key Contribution to the Host Airway-Pathogen Interplay. ACS Infect Dis 6:406–421.

34. Jiang D, Tikhomirova A, Kidd SP. 2016. Haemophilus influenzae strains possess variations in the global transcriptional profile in response to oxygen levels and this influences sensitivity to environmental stresses. Res Microbiol 167:13–9.

35. Muda NM, Nasreen M, Dhouib R, Hosmer J, Hill J, Mahawar M, Schirra HJ, McEwan AG, Kappler U. 2019. Metabolic analyses reveal common adaptations in two invasive Haemophilus influenzae strains. Pathog Dis 77.

36. Pifer ML, Smith HO. 1985. Processing of donor DNA during Haemophilus influenzae transformation: analysis using a model plasmid system. Proc Natl Acad Sci U S A 82:3731–5.

37. MacFadyen LP, Chen D, Vo HC, Liao D, Sinotte R, Redfield RJ. 2001. Competence development by Haemophilus influenzae is regulated by the availability of nucleic acid precursors. Mol Microbiol 40:700–7.

38. Redfield RJ. 1993. Genes for breakfast: the have-your-cake-and-eat-it-too of bacterial transformation. J Hered 84:400–4.

39. Macfadyen LP, Dorocicz IR, Reizer J, Saier MH, Redfield RJ. 1996. Regulation of competence development and sugar utilization in Haemophilus influenzae Rd by a phosphoenolpyruvate:fructose phosphotransferase system. Mol Microbiol 21:941–52.

40. Chandler MS. 1992. The gene encoding cAMP receptor protein is required for competence development in Haemophilus influenzae Rd. Proc Natl Acad Sci U S A 89:1626–30.

41. Dorocicz IR, Williams PM, Redfield RJ. 1993. The Haemophilus influenzae adenylate cyclase gene: cloning, sequence, and essential role in competence. J Bacteriol 175:7142–9.

42. Rubin BK. 2007. Mucus structure and properties in cystic fibrosis. Paediatr Respir Rev 8:4–7.

43. Matthews LW, Spector S, Lemm J, Potter JL. 1963. Studies on pulmonary secretions. I. The over-all chemical composition of pulmonary secretions from patients with cystic fibrosis, bronchiectasis, and laryngectomy. Am Rev Respir Dis 88:199–204.

44. Herriott RM, Meyer EM, Vogt M. 1970. Defined nongrowth media for stage II development of competence in Haemophilus influenzae. J Bacteriol 101:517–24.

45. Wong SMS, Akerley BJ. 2012. Genome-scale approaches to identify genes essential for Haemophilus influenzae pathogenesis. Front Cell Infect Microbiol 2:23.

46. McFarland WC, Stocker BA. 1987. Effect of different purine auxotrophic mutations on mouse-virulence of a Vi-positive strain of Salmonella dublin and of two strains of Salmonella typhimurium. Microb Pathog 3:129–41.

47. Wang J, Mushegian A, Lory S, Jin S. 1996. Large-scale isolation of candidate virulence genes of Pseudomonas aeruginosa by in vivo selection. Proc Natl Acad Sci U S A 93:10434–9.

48. Yee R, Cui P, Shi W, Feng J, Zhang Y. 2015. Genetic Screen Reveals the Role of Purine Metabolism in Staphylococcus aureus Persistence to Rifampicin. Antibiotics (Basel) 4:627–42.

49. Berney M, Berney-Meyer L. 2017. Mycobacterium tuberculosis in the Face of Host-Imposed Nutrient Limitation. Microbiol Spectr 5.

50. Polissi A, Pontiggia A, Feger G, Altieri M, Mottl H, Ferrari L, Simon D. 1998. Large-scale identification of virulence genes from Streptococcus pneumoniae. Infect Immun 66:5620–9.

51. Lau GW, Haataja S, Lonetto M, Kensit SE, Marra A, Bryant AP, McDevitt D, Morrison DA, Holden DW. 2001. A functional genomic analysis of type 3 Streptococcus pneumoniae virulence. Mol Microbiol 40:555–71.

52. Hava DL, Camilli A. 2002. Large-scale identification of serotype 4 Streptococcus pneumoniae virulence factors. Mol Microbiol 45:1389–406.

53. Ge X, Kitten T, Chen Z, Lee SP, Munro CL, Xu P. 2008. Identification of Streptococcus sanguinis genes required for biofilm formation and examination of their role in endocarditis virulence. Infect Immun 76:2551–9.

54. Kim JK, Jang HA, Won YJ, Kikuchi Y, Heum Han S, Kim C-H, Nikoh N, Fukatsu T, Lee BL. 2014. Purine biosynthesis-deficient Burkholderia mutants are incapable of symbiotic accommodation in the stinkbug. ISME J 8:552–563.

55. Gawronski JD, Wong SMS, Giannoukos G, Ward D v, Akerley BJ. 2009. Tracking insertion mutants within libraries by deep sequencing and a genome-wide screen for Haemophilus genes required in the lung. Proc Natl Acad Sci U S A 106:16422–7.

56. Ganz T. 2018. Iron and infection. Int J Hematol 107:7–15.

57. Archibald FS, DeVoe IW. 1979. Removal of iron from human transferrin by Neisseria meningitidis. FEMS Microbiol Lett 6:159–162.

58. McKenna WR, Mickelsen PA, Sparling PF, Dyer DW. 1988. Iron uptake from lactoferrin and transferrin by Neisseria gonorrhoeae. Infect Immun 56:785–91.

59. Gray-Owen SD, Loosmore S, Schryvers AB. 1995. Identification and characterization of genes encoding the human transferrin-binding proteins from Haemophilus influenzae. Infect Immun 63:1201–10.

60. Parker Siburt CJ, Mietzner TA, Crumbliss AL. 2012. FbpA — A bacterial transferrin with more to offer. Biochimica et Biophysica Acta (BBA) - General Subjects 1820:379–392.

61. Whitby PW, Seale TW, VanWagoner TM, Morton DJ, Stull TL. 2009. The iron/heme regulated genes of Haemophilus influenzae: comparative transcriptional profiling as a tool to define the species core modulon. BMC Genomics 10:6.

62. Morton DJ, Musser JM, Stull TL. 1993. Expression of the Haemophilus influenzae transferrin receptor is repressible by hemin but not elemental iron alone. Infect Immun 61:4033–7.

63. Zhong Q, Kobe B, Kappler U. 2020. Molybdenum Enzymes and How They Support Virulence in Pathogenic Bacteria. Front Microbiol 11:615860.

64. Leimkühler S, Iobbi-Nivol C. 2016. Bacterial molybdoenzymes: old enzymes for new purposes. FEMS Microbiol Rev 40:1–18.

65. Hille R, Hall J, Basu P. 2014. The mononuclear molybdenum enzymes. Chem Rev 114:3963–4038.

66. Imperial J, Ugalde RA, Shah VK, Brill WJ. 1985. Mol-mutants of Klebsiella pneumoniae requiring high levels of molybdate for nitrogenase activity. J Bacteriol 163:1285–7.

67. Maia LB, Moura JJG, Moura I. 2015. Molybdenum and tungsten-dependent formate dehydrogenases. J Biol Inorg Chem 20:287–309.

68. Schroeder HA, Balassa JJ, Tipton IH. 1970. Essential trace metals in man: molybdenum. J Chronic Dis 23:481–99.

69. Evans SS, Repasky EA, Fisher DT. 2015. Fever and the thermal regulation of immunity: the immune system feels the heat. Nat Rev Immunol 15:335–49.

70. Small PM, Täuber MG, Hackbarth CJ, Sande MA. 1986. Influence of body temperature on bacterial growth rates in experimental pneumococcal meningitis in rabbits. Infect Immun 52:484–7.

71. Green MH, Vermeulen CW. 1994. Fever and the control of gram-negative bacteria. Res Microbiol 145:269–72.

72. Smoot LM, Smoot JC, Graham MR, Somerville GA, Sturdevant DE, Migliaccio CA, Sylva GL, Musser JM. 2001. Global differential gene expression in response to growth temperature alteration in group A Streptococcus. Proc Natl Acad Sci U S A 98:10416– 21.

73. Schortgen F. 2012. Fever in sepsis. Minerva Anestesiol 78:1254–64.

74. Kappler U, Nasreen M, McEwan A. 2019. New insights into the molecular physiology of sulfoxide reduction in bacteria. Adv Microb Physiol 75:1–51.

75. Nasreen M, Dhouib R, Hosmer J, Wijesinghe HGS, Fletcher A, Mahawar M, Essilfie A-T, Blackall PJ, McEwan AG, Kappler U. 2020. Peptide Methionine Sulfoxide Reductase from Haemophilus influenzae Is Required for Protection against HOCl and Affects the Host Response to Infection. ACS Infect Dis 6:1928–1939.

76. Nasreen M, Nair RP, McEwan AG, Kappler U. 2022. The Peptide Methionine Sulfoxide Reductase (MsrAB) of Haemophilus influenzae Repairs Oxidatively Damaged Outer Membrane and Periplasmic Proteins Involved in Nutrient Acquisition and Virulence. Antioxidants (Basel) 11.

77. Dhouib R, Nasreen M, Othman DSMP, Ellis D, Lee S, Essilfie A-T, Hansbro PM, McEwan AG, Kappler U. 2021. The DmsABC Sulfoxide Reductase Supports Virulence in Non-typeable Haemophilus influenzae. Front Microbiol 12:686833.

